# DNA damage response at telomeres boosts the transcription of SARS-CoV-2 receptor ACE2 during aging

**DOI:** 10.1101/2021.06.09.447484

**Authors:** Sara Sepe, Francesca Rossiello, Valeria Cancila, Fabio Iannelli, Valentina Matti, Giada Cicio, Matteo Cabrini, Eugenia Marinelli, Busola Alabi, Alessia di Lillo, Arianna Di Napoli, Jerry W. Shay, Claudio Tripodo, Fabrizio d’Adda di Fagagna

## Abstract

The severe acute respiratory syndrome coronavirus 2 (SARS-CoV-2) infection is known to be more common in the elderly, who show also more severe symptoms and a higher risk of hospitalization and death. Here we show that the expression of the Angiotensin Converting Enzyme 2 (ACE2), the SARS-CoV2 cell receptor, increases during aging in mouse and human lungs, and following telomere shortening or dysfunction in mammalian cells and in mouse models. This increase is regulated at the transcription level, and *Ace2* promoter activity is DNA damage response (DDR)-dependent. Indeed, ATM inhibition or the selective inhibition of telomeric DDR, through the use of antisense oligonucleotides, prevents *Ace2* upregulation following telomere damage, in cultured cells and in mice.

We propose that during aging telomeric shortening, by triggering DDR activation, causes the upregulation of ACE2, the SARS-CoV2 cell receptor, thus making the elderly likely more susceptible to the infection.

## Introduction

The severe acute respiratory syndrome coronavirus 2 (SARS-CoV-2) is responsible for the recent coronavirus disease 2019 pandemic. The clinical course of the disease is variable, ranging from asymptomatic infection to multi-organ failure and death, with a fatality rate of ~5%. The severity of the infection has been correlated with the age of patients, suggesting that aging is an important risk factor influencing the outcome (Verity *et al*, 2020). The reasons for the development of severe symptoms in the elderly compared to young individuals are still under intense investigation. The expression of angiotensin-converting enzyme 2 (ACE2), which works as the host functional receptor for SARS-CoV-2 (Yan R *et al*, 2020)(Lan *et al*, 2020), has been positively related to patients’ age (Beyerstedt *et al*, 2021), although some reports question the effective increase of ACE2 levels during aging (Xudong *et al*, 2006).

## Results and Discussion

In order to better clarify the modulation of the ACE2 expression during aging, and the potential molecular mechanisms underlying it, we studied the expression of ACE2 in mouse and human lungs at different ages.

We observed an increase of *Ace2* mRNA levels (Fig 1A) in lungs of aged mice (22-24 months) compared to young mice (2-3 months) as detected by RT-qPCR. Immunohistochemistry (IHC) analyses of the same lungs confirmed and extended this observation to the protein level. ACE2 protein was detected in both pneumocytic and stromal/inflammatory elements in the lungs of old mice (Fig 1B). Quantitative analysis of IHC labeling showed a significant increase in terms of density of expression in old mice, as compared with young mice (Fig 1B).

**Figure 1.**
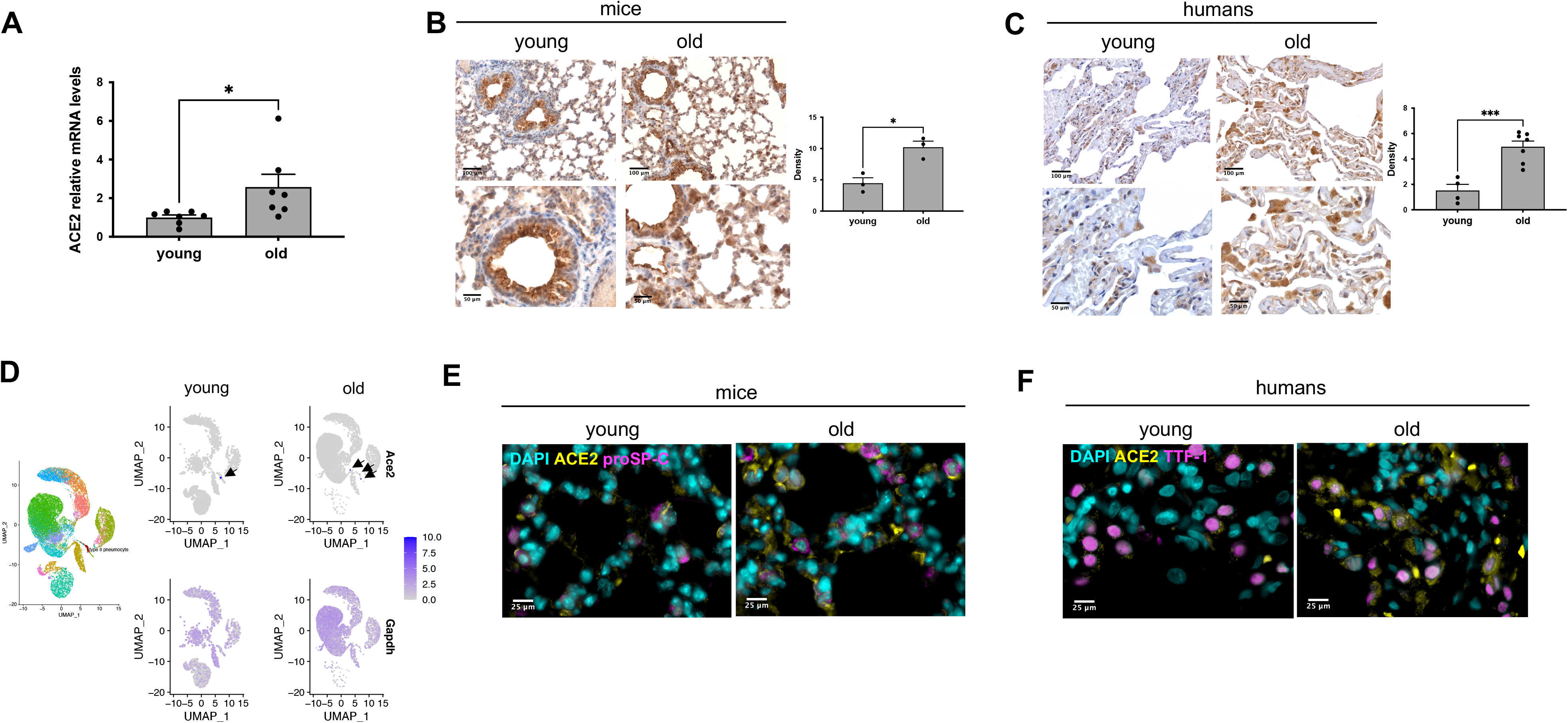
ACE2 expression increases during ageing in mouse and human lungs. **A** RT-qPCR detection of *Ace2* mRNA expression levels in lungs from young (2-3 months) and old (22-24 months) mice (n = 7 mice per group). Error bars represent the s.e.m. *P<0.05. **B** Representative microphotographs and quantitative analyses of ACE2 immunohistochemical staining in lungs from young (2 months) and old (22 months) mice (n=3 mice/group). Original magnification x200 and x400. Scale bar, 100 and 50 μm. Error bars represent the s.e.m. *P<0.05 **C** Representative microphotographs and quantitative analyses of ACE2 immunohistochemical staining in lung parenchyma of young (20-35 years old) and old (60-80 years old) humans (n = 3/5 individuals per group). Original magnification x200 and x400. Scale bar, 100 and 50 μm. Error bars represent the s.e.m. ***P<0.001 **D** Dimension reduction plot showing *Ace2* and *Gadph* expression levels according to age in type II pneumocytes in single-cell transcriptomic data from aging tissues in the mouse lung from Tabula Muris Senis. Left panel show cells belonging to the type II pneumocyte cluster as red circles with black outline. The right 4 panels show *Ace2* and *Gadph* expression levels respectively at 3 (young) and 30 (old) months, according to an increasing blue scale. **E** Double-marker immunofluorescence for ACE2 and pro-SP-C in lungs from young and old mice detailing the presence of ACE2 in type II pneumocytic pro-SP-C positive elements (n=3 mice per group). Original magnifications, x630. Scale bar, 25 μm. **F** Double-marker immunofluorescence for ACE2 and TTF-1 in lungs parenchyma from young and old human tissues detailing the expression of ACE2 in type II pneumocytic TTF-1 positve cells (n = 3/5 individuals per group). Original magnifications, x630. Scale bar, 25 μm.

Prompted by the detection of ACE2 different expression levels in lungs from young and old mice, we extended our analyses to human lung tissue samples by comparing ACE2 immunostaining in histologically normal lung parenchyma of young (20-35 years old) subjects and old (60-80 years old) subjects. ACE2 protein expression levels in the lung parenchyma of old subjects proved to be higher as compared with that of young subjects, as shown and quantified in Fig 1C.

Emerging evidence suggests that the severity of SARS-CoV-2 infection correlates with high rates of alveolar epithelial type II (ATII) cells infection (Yee *et al*, 2020). We took advantage of the single-cell transcriptomic mouse atlas spanning different ages (Almanzar *et al*, 2020) and focused on the expression data of lungs. We observed that, while housekeeping genes such as *Gapdh* are widely expressed in almost all cell types, *Ace2* showed an ATII pneumocytes-preferential expression that increased upon aging (Fig 1D and EV1A). These conclusions are consistent with similar analyses performed on non-human primates (Ziegler *et al*, 2020).

To confirm that ACE2-expressing elements were actually ATII cells, we performed double-marker immunofluorescence for ACE2 and pro-SP-C in mice or TTF-1 in human lung samples of different ages. Immunofluorescence revealed that ACE2 was predominantly expressed in ATII pneumocytes and it increased with aging (Figs 1E-F). These results indicate that ACE2 expression is upregulated with age in lungs, thus likely favoring infection by SARS-CoV-2 and causing more severe symptoms.

Aging is associated with telomere shortening and damage in several tissues (Vaiserman & Krasnienkov, 2021)(Demanelis *et al*, 2020)(Fumagalli *et al*, 2012)(Hewitt *et al*, 2012). To test if telomere shortening can modulate *ACE2* expression, we monitored its mRNA levels in human fibroblasts (BJ) and human bronchial epithelial cells (HBECs) at different population doublings since both cell types do not have telomere maintenance mechanisms and are characterized by progressive telomere shortening upon proliferation (Huffman *et al*, 2000)(Peters-Hall *et al*, 2018). Both late passage BJ and HBEC had increased levels of *ACE2* mRNA, compared to cells at early passages (Fig 2A-B). Recently, a shorter *ACE2* splicing isoform has been described and proposed to be upregulated by interferon response upon infection by different viruses (but not upon SARS-Cov-2). Since the shorter splice isoform lacks the extracellular domain, that is used as SARS-CoV-2 binding site, it is not relevant for SARS-CoV-2 infection (Blume *et al*, 2021). The observed increase in late passages BJ cells was not due to this short isoform, since primers specific for the long isoform (see materials and methods section) detected a similar increase in RT-qPCR assays (Fig EV1B), while the short isoform was below detection levels (data not shown).

**Figure 2.**
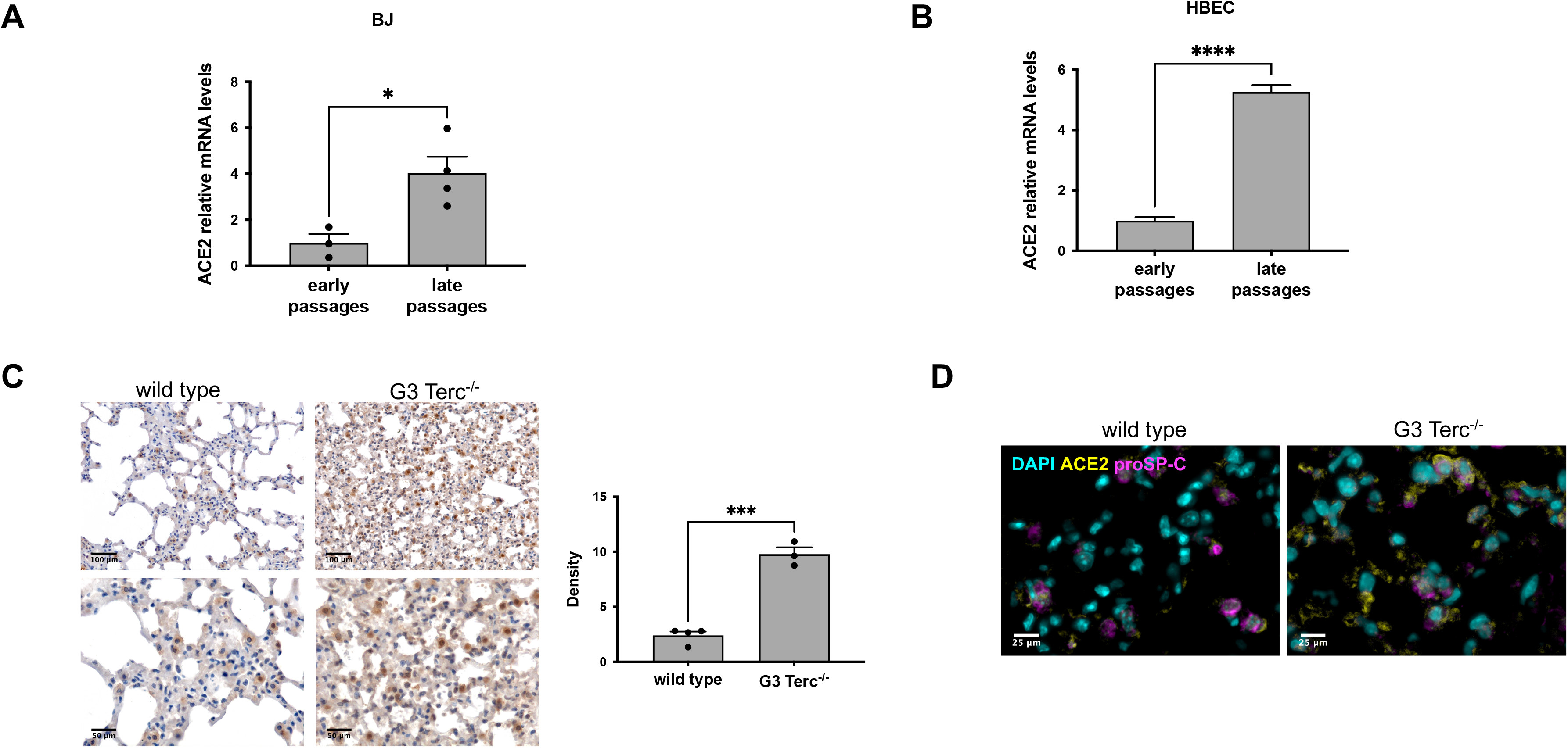
ACE2 levels increase upon telomere shortening in human cells and in lung of G3 Terc^−/−^ mice. **A** RT-qPCR detection of *ACE2* mRNA expression levels in early passages (PD 34-37) and late passages (PD 61-64) human normal fibroblasts (BJ) (n = 3-4 replicates per group). Error bars represent the s.e.m. *P<0.05 **B** ddPCR detection of *ACE2* mRNA expression levels in early passages (PD 22.37) and late passages (PD 140) human bronchial epithelial cells (HBEC) (n = 6 replicates per group). Error bars represent the s.e.m. ****P<0.0001. **C** Representative images and quantitative analyses of ACE2 immunohistochemical staining in lungs of age-matched wild type and G3 Terc^−/−^ mice (n = 3/4 mice per group). Original magnification x200 and x400. Scale bar, 100 and 50 μm. Error bars represent the s.e.m. ***P<0.001. **D** Double-marker immunofluorescence for ACE2 and pro-SP-C in lungs of age-matched wild type and G3 Terc^−/−^ mice (n = 3-4 mice per group). Original magnifications, x630. Scale bar, 25 μm.

We extended our studies to a mouse model lacking the RNA component of telomerase (Terc^−/−^), which at late generations recapitulates several features of human aging in different tissues (Lee *et al*, 1998)(Rudolph *et al*, 1999)(Giorgio *et al*, 2016), including lungs (Piñeiro-Hermida *et al*, 2020). When we analyzed the expression level and the distribution of ACE2 protein in the lungs of the third generation (G3) of Terc^−/−^ mice at 12 months of age and in age- and sex-matched wild type animals, we observed a consistent increase of ACE2 expression in G3 Terc^−/−^ mice when compared to wild type counterparts (Fig 2C). Immunofluorescence imaging with specific cell markers demonstrated that ACE2 increases mainly in ATII pneumocytes expressing pro-SP-C (Fig 2D).

When telomeres become critically short, they activate the DNA damage response (DDR) pathway (D’Adda Di Fagagna *et al*, 2003)(Herbig *et al*, 2004). To test if telomeric DDR is sufficient to increase *ACE2* mRNA levels, we used two mammalian cell systems, which allow the activation of the DDR specifically at telomeres, in the absence of telomere shortening. TRF2 is a component of the shelterin complex and prevents telomeric DNA from being recognized as a DNA lesion and activate a DDR (Palm & De Lange, 2008).

We used *Trf2* conditional knockout mouse embryonic fibroblasts (MEFs Trf2^F/F^) carrying a Cre recombinase (Rosa26-CreERT2) inducible by 4-hydroxytamoxifen (4OHT) (Okamoto *et al*, 2013) and HeLa cells with a doxycycline-regulated expression of a short hairpin against TRF2 (HeLa shTRF2) to knockdown TRF2 expression (Grolimund *et al*, 2013).

TRF2 knockout or knockdown led to telomeric DDR activation as shown by foci of DDR markers at telomeres and increased levels of telomeric damage-induced long non-coding RNA (tdilncRNA) (Aguado *et al*, 2019)(Rossiello *et al*, 2017) in both cell lines (Figs 3A,C and EV2A). Here, we observed a correlation between DDR increase and *ACE2* mRNA levels (Figs 3B,D). Interestingly, DDR activation following ionizing radiation (IR) in the same cell lines (Figs EV2B,D) drove a similar increase of *ACE2* mRNA levels (Figs EV2C,E), indicating that while telomeric DDR activation is the most likely event increasing ACE2 levels during physiological aging, additional events activating DDR could also contribute. To verify the effects of telomeric DDR *in vivo*, we next used an inducible Trf2 knockout mouse model (Trf2^fl/fl^ mice) (Rossiello *et al*, 2017) in which, following tamoxifen administration, *Trf2* expression is lost, leading to DDR activation at telomeres as shown by the accumulation of foci of 53BP1, a robust marker of DDR activation (Fig 3E). Parallel to DDR activation, we observed an increase of *Ace2* mRNA levels in mouse tissues upon tamoxifen administration (Fig 3F), demonstrating that also *in vivo* the activation of DDR at telomere leads to increased *Ace2* expression.

**Figure 3.**
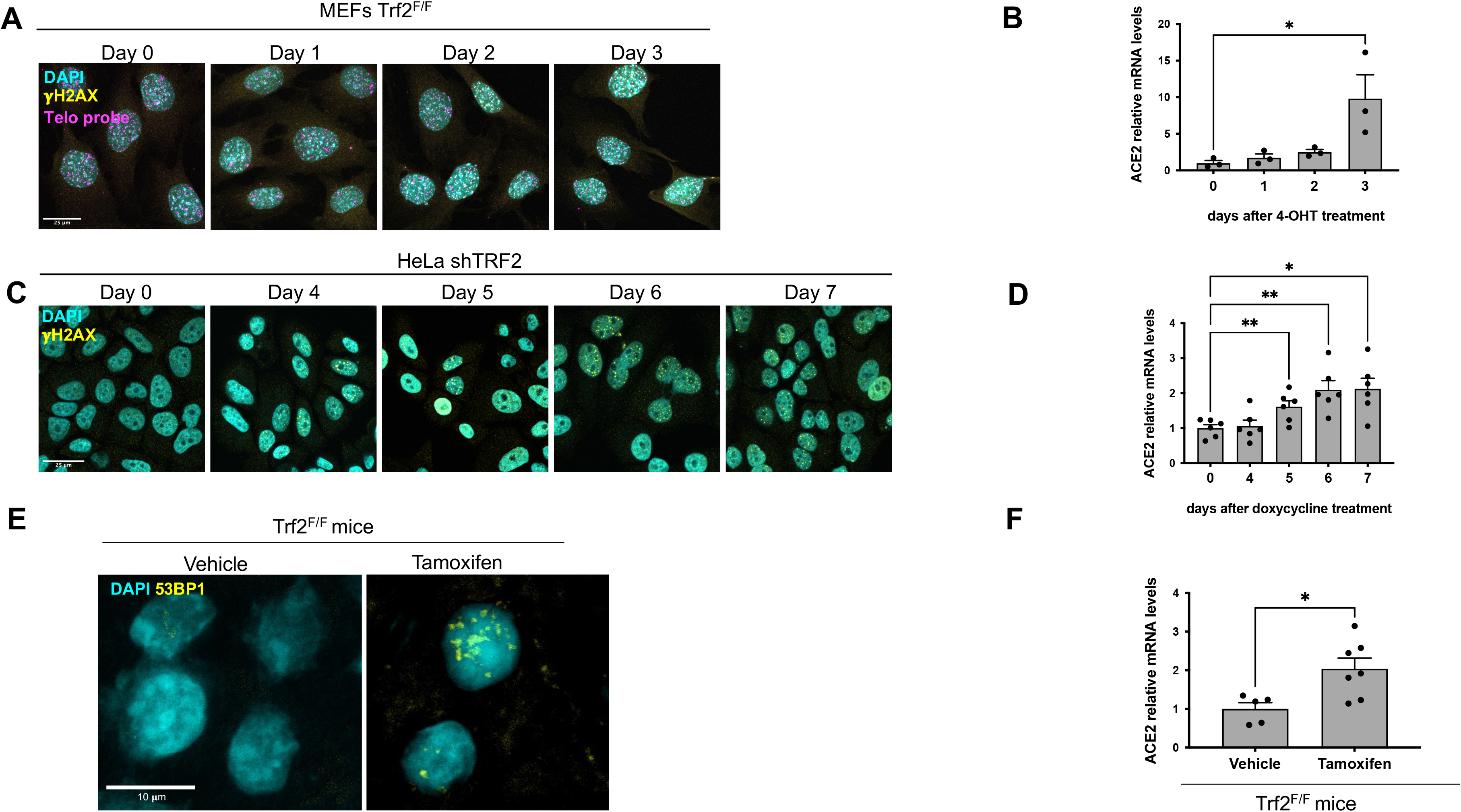
DDR activation induces ACE2 expression in cultured cells and *in vivo*. **A** ImmunoFISH showing co-localization of γH2AX foci and the telomeric probe in MEFs Trf2^F/F^ at the indicated time points following 4OHT treatment and consequent TRF2 knockout. Scale bar, 25 μm. **B** RT-qPCR detection of *Ace2* mRNA expression levels in MEFs Trf2^F/F^ treated as in A (n = 3 independent experiments). Error bars represent the s.e.m. *P<0.05 **C** Immunofluorescence showing γH2AX foci in HeLa shTRF2 cells at the indicated time points following doxycycline treatment and consequent TRF2 knockdown. **D** RT-qPCR detection of *ACE2* mRNA expression levels in HeLa shTRF2 cells treated as in C (n = 6 independent experiments). Error bars represent the s.e.m. *P<0.05, **P<0.01 **E** Representative immunofluorescence images of 53BP1 staining in liver from Trf2^F/F^ mice treated with tamoxifen (to induce TRF2 loss and telomere uncapping) or vehicle. **F** RT-qPCR detection of *Ace2* mRNA expression levels in livers of mice treated as in E (n = 5-7 mice/group). Error bars represent the s.e.m. *P<0.05.

Taken together, these results in human and mouse cell lines and in mouse tissues indicate a role for the activated DDR pathways in the increase of ACE2 levels.

The increase of mRNA levels can be caused by a regulated process at the *ACE2* transcriptional promoter level. In order to determine whether the *ACE2* promoter could respond to DDR activation, we first performed an *in silico* analysis to identify the transcription factors DNA binding motifs significantly enriched in the promoter region of *ACE2*. Gene set enrichment analysis of the top 100 transcription factors potentially associated with the *ACE2* promoter, revealed an over-representation of pathways possibly related to the DNA damage response (Fig 4A).

**Figure 4.**
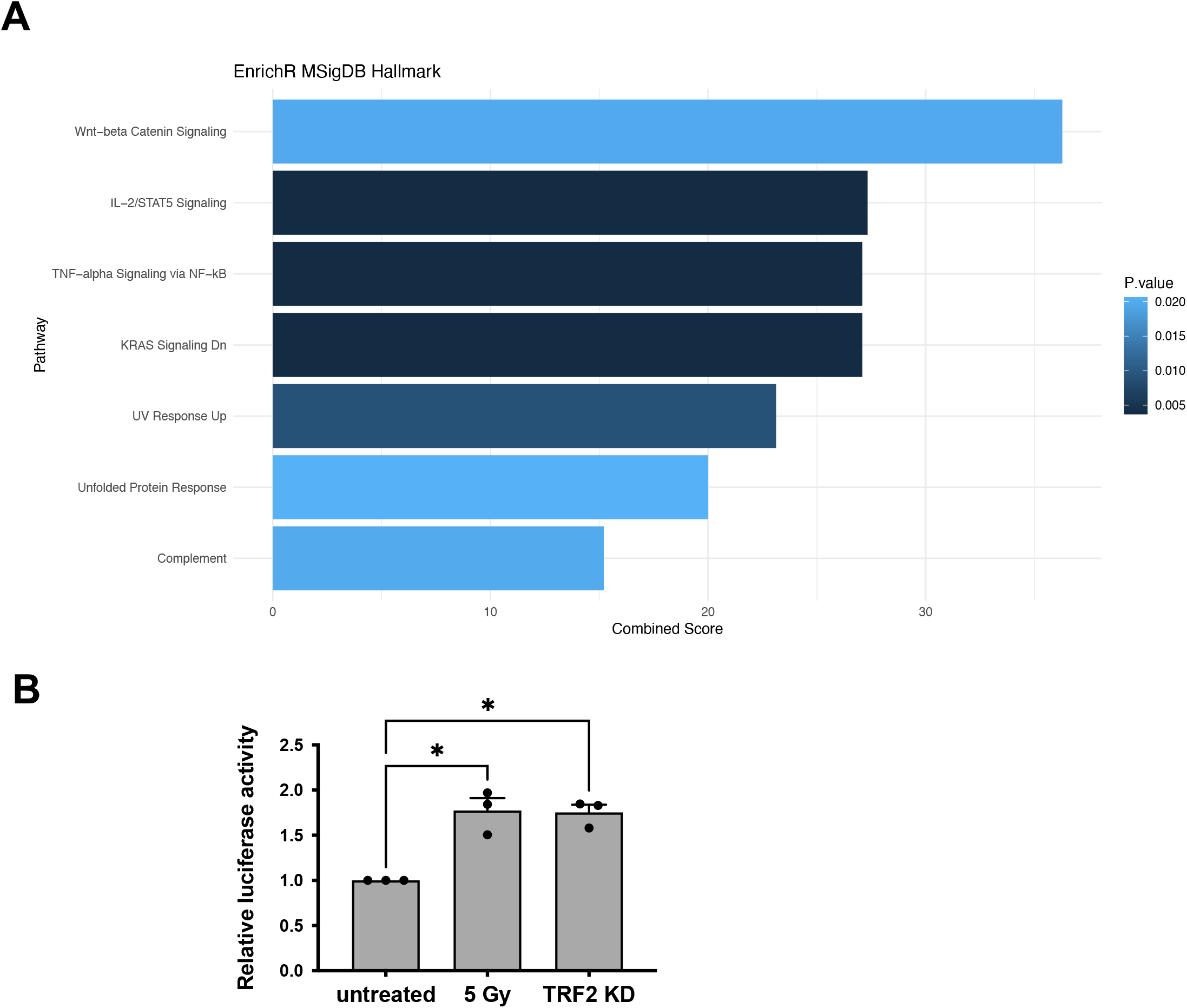
DDR activation mediates transcriptional upregulation of *ACE2* gene. **A** Gene set enrichment analysis showing significantly enriched pathways (P < 0.05) from the MSigDB_Hallmark gene-set library, bars are colored with an increasing blue scale according to p value. Combined score: combined value of both the p-value and z-score. **B** Relative luciferase activity in HeLa shTRF2 following ionizing radiation (5 Gy) or TRF2 knockdown upon doxycycline-induced shTRF2 expression (n=3 independent experiments). Error bars represent the s.e.m. *P<0.05.

To experimentally demonstrate that the *ACE2* promoter responds to the activation of the DDR pathways, we transfected a plasmid carrying the luciferase gene as a reporter under the control of human *ACE2* promoter in HeLa shTRF2 cells and we induced telomeric DDR by TRF2 knockdown. Importantly, we observed an increase of luciferase signal upon telomere dysfunction. A similar transcriptional activation was also observed also upon exposure of uninduced HeLa cells to IR (Fig 4B).

To demonstrate that DDR pathway activation is directly responsible for the observed increase in *Ace2* mRNA levels, we treated MEFs Trf2^F/F^ with the ATM kinase inhibitor KU-60019 (ATMi) while inducing Trf2 knockout by 4OHT treatment. While, as expected, this treatment prevented the DDR foci formation (Fig 5A), it also significantly reduced *Ace2* mRNA levels increase (Fig 5B), demonstrating ATM kinase activity involvement in *Ace2* transcripts levels.

**Figure 5.**
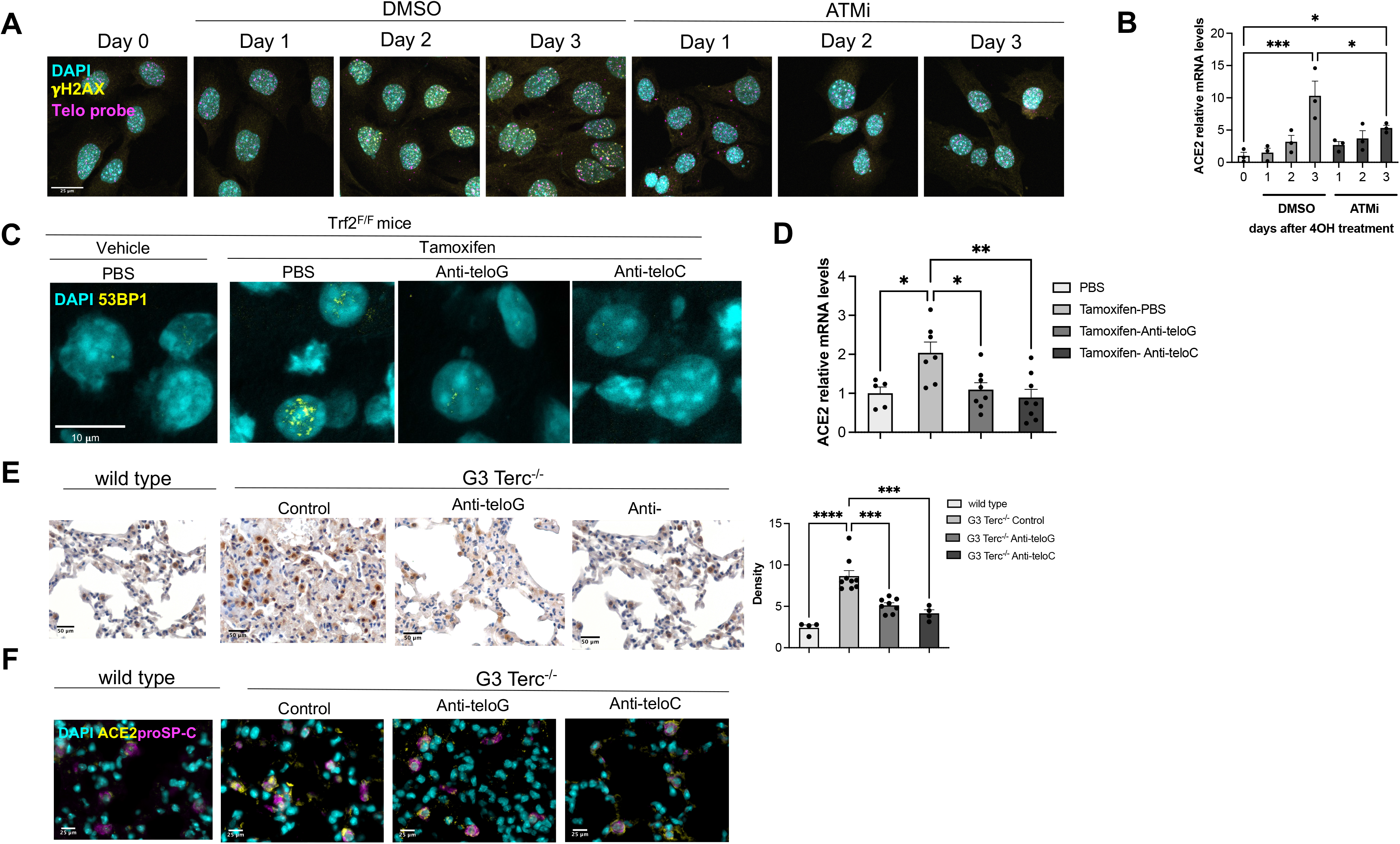
Selective inhibition of telomeric DDR decreases ACE2 expression in cultured cells and *in vivo*. **A** ImmunoFISH showing co-localization of γH2AX foci and the telomeric probe in MEFs Trf2^F/F^ at the indicated time points following 4OHT treatment and consequent TRF2 knockout and treated with DSMO or ATMi (KU-60019, 10 μM). Scale bar, 25 μm. **B** RT-qPCR detection of *Ace2* mRNA expression levels in MEFs Trf2^F/F^ treated as in A (n=3 independent experiments). Error bars represent the s.e.m. *P<0.05, ***P<0.001 **C** Representative immunofluorescence images of 53BP1 staining in liver from Trf2^F/F^ mice treated with tamoxifen (to induce telomere uncapping) or vehicle and injected with the indicated ASOs or PBS as control. Scale bar, 10 μm. **D** RT-qPCR detection of *Ace2* mRNA levels in livers of mice treated as in C (n= 5-8 mice per group). Error bars represent the s.e.m. *P<0.05, **P<0.01 **E** Representative microphotographs and quantitative analyses of ACE2 immunohistochemical staining in lungs of age-matched wild type and G3 Terc^−/−^ mice, treated with the indicated ASOs (n = 4/8 mice per group). Original magnification x200 and x400. Scale bar, 100 and 50 μm. Error bars represent the s.e.m. ***P<0.001, ****P<0,0001 **F** Double-marker immunofluorescence for ACE2 and pro-SP-C in lungs of age-matched wild type and G3 Terc^−/−^ mice, treated with the indicated ASOs (n = 4-8 mice/group). Original magnifications, x630. Scale bar, 25 μm.

In order to investigate if inhibition of DDR could affect the expression of ACE2 also *in vivo*, we took advantage of telomeric antisense oligonucleotides (tASOs), which allow the specific inhibition of telomeric DDR by targeting tdilncRNA generated at dysfunctional telomeres and necessary for full DDR activation at telomeres (Michelini *et al*, 2017) (Rossiello *et al*, 2017)(Aguado *et al*, 2019). To test the impact of selective telomeric DDR inhibition on ACE2 expression, following tamoxifen-induced *Trf2* loss, mice were treated with a systemic dose of anti-teloG or anti-teloC ASOs, or phosphate-buffered saline (PBS) as negative control, delivered by intraperitoneal injection. Four days later, when telomeric DDR reached maximal activation in this model, mice were sacrificed and the liver was analysed. Both tASO treatments were highly effective, as demonstrated by decreased number of 53BP1 foci (Fig 5C) as previously shown (Rossiello *et al*, 2017), and led to *Ace2* mRNA levels downregulation *in vivo* (Fig 5D). Stimulated by this result, we then extended our studies to the G3 Terc^−/−^ mouse model of telomere shortening and dysfunction. We treated 3 months old G3 Terc^−/−^ mice with anti-teloG or anti-teloC ASOs or an ASO with an unrelated sequence (Control) by intraperitoneal injection and we sacrificed them at 12 months of age to collect their lungs.

Quantitative immunohistochemical analyses of ACE2 expression revealed high levels of the protein in Control G3 Terc^−/−^ mice, compared to age- and sex-matched wild type animals. Such an expression was significantly dampened by the treatment with either tASOs (Fig 5E), which impacted on ACE2 expression mainly in pro-SP-C-expressing pneumocytic elements, as shown by double-marker immunofluorescence (Fig 5F). These experiments also indicate that, rather than telomeric shortening *per se*, it is the ensuing DDR activation that is responsible for ACE2 regulation.

Taken together, these results clearly demonstrate that the expression of ACE2, the SARS-CoV-2 receptor, is directly modulated by the activation of the DDR pathway at the transcriptional level, and that telomere dysfunction is a physiological event able to engage the DDR pathways modulating ACE2 levels. Our results also demonstrate that tASOs are agents effective in controlling ACE2 levels *in vivo* in mouse tissues, including the lung. In perspective, as ASOs are established medicines approved for a number of disorders (Crooke *et al*, 2021) and as telomeric DDR increases with age in normal and pathological conditions (Rossiello et al, Nature Cell Biology, under submission) we suggest that, upon additional validation, tASOs can have a potential as therapeutic agents to reduce susceptibility to SARS-CoV-2 infection.

## Materials and Methods

### Cells and treatments

Foreskin fibroblast BJ cells (The American Type Culture Collection) were grown in MEM supplemented with 10% fetal bovine serum, 1% glutamine, 10 mM non-essential amino acids and 1 mM sodium pyruvate.

Primary Human bronchial epithelial cells (HBECs) were co-cultured with irradiated 3T3 J2 feeder cells with ROCK inhibitor and 2% O_2_ (ROCKi conditions) as described previously (Peters-Hall *et al*, 2018).

Rosa26-CreERT2 Trf2^F/F^ MEFs (Okamoto *et al*, 2013), a kind gift from Eros Lazzerini Denchi (National Cancer Institute, NIH, Bethesda, USA), were grown in DMEM supplemented with 10% fetal bovine serum and 1% glutamine; for CreER activation, cells were treated with 600nM 4 hydroxytamoxifen (4OHT, H7904, Sigma-Aldrich).

HeLa inducible shTRF2 cells (Grolimund *et al*, 2013), a gift from Joachim Lingner (ISREC-EPFL, Lausanne, Switzerland), expressing the tTR-KRAB construct, the dsRED marker and the doxycycline-inducible shTRF2 and GFP marker, were grown in DMEM supplemented with 10% Tet System Approved fetal bovine serum and 1% glutamine; for shTRF2 and GFP induction, cells were treated with doxycycline (1 μg ml^−1^).

All cell lines used in this study were grown under standard cell culture conditions (37°C, 5% CO_2_) and were tested negative for mycoplasma contaminations.

Ionizing radiation was induced by a high-voltage X-ray-generator tube (Faxitron X-Ray Corporation).

ATM kinase inhibitor KU-60019 (S1570 Selleckchem), or dimethylsulphoxide as negative control, was used at 10 μM concentration.

### Animals and treatments

Experiments involving animals have been done in accordance with the Italian Laws (D.lgs. 26/2014), which enforce Directive 2010/63/EU (Directive 2010/63/EU of the European Parliament and of the Council of 22 September 2010 on the protection of animals used for scientific purposes). Accordingly, the project has been authorized by the Italian Competent Authority (Ministry of Health).

For the analysis of young and old mice, C57BL/6 J mice were purchased from the Charles River Laboratories. Young animals were sacrificed at 2-3 months of age, while old animals at 22-24 months of age. After collection, lungs were snap frozen for RNA extraction and formalin-fixed and paraffin embedded for histological analysis.

Terc mice (B6.Cg-Terctm1Rdp/J Stock No: 004132 | mTR^−/−^)(Blasco *et al*, 1997) were purchased by Jackson Laboratory. Terc^+/−^ mice were intercrossed to generate first-generation (G1) homozygous Terc^−/−^ knockout mice. Second-generation (G2) Terc^−/−^ mice were generated by successive breeding of G1 Terc^−/−^ and then G3 Terc^−/−^ mice by crosses between G2 Terc^−/−^ mice. Finally, wild type C57BL/6J and G3 Tert^−/−^ mice were used for the experiments. The 8-10 weeks-old mice were injected intraperitoneally (i.p.) with a Control, ASO anti-teloG and ASO anti-teloC at a final concentration of 15 mg/kg twice a week for four weeks. The animals were sacrificed at 12 months of age. The lungs were collected and part of it was snap frozen for RNA extraction and part was washed in PBS and collected for fixation in 10% neutral buffered formalin overnight, washed in water and paraffin-embedded for histological analysis.

Rosa26-CreERT mice (Jackson Laboratory) and TRF2 conditional knockout mice (Celli & de Lange, 2005) and mice carrying a p53 conditional allele (Jackson Laboratory) were crossed to generate Trf2/p53/Rosa26 mice. Mice were maintained in 129/c57Bl6 genetic background. All mice were bred and maintained under pathogen-free condition at the Scripps Research Institute and were handled according to Institutional Animal Care and Use Committee guidelines. The animals were provided by Eros Lazzerini Denchi (National Cancer Institute, NIH, Bethesda, USA). To activate CreER, 8 to 10-weeks-old mice were injected intraperitoneally (i.p.) with tamoxifen dissolved in sunflower oil or with vehicle at a final concentration of 75 mg/kg. After 24 hours Vehicle (PBS), ASO anti-teloG and anti-teloC dissolved in PBS were administrated by i.p. at a concentration of 15 mg/kg. Mice were sacrificed after five days post tamoxifen injection. Tissues were collected and frozen in dry ice and embedded in OCT tissue TEC (Sakura).

### Human samples

Human lung tissue samples were collected from the archives of the Department of Clinical and Molecular Medicine, Pathology Unit, Sant’Andrea Hospital, Sapienza University, Rome, Italy. Samples were relative to histologically normal parenchyma adjacent to either foci of subpleural emphysema (young subjects) or non-small cell lung cancer (old subjects). The samples were collected and handled according to the Helsinki Declaration. Patients provided written informed consent to the use of their tissue samples for research purposes (Ethical Approval SA250).

### Antisense oligonucleotides (ASOs) sequences

The locked nucleic acid-modified oligonucleotides with a fully phosphorothioate backbone were produced by Qiagen as described (Rossiello *et al*, 2017).

Sequences were as follows (5’–3’ orientation):

Control: ACTGATAGGGAGTGGTAAACT
anti-teloG: CCCTAACCCTAACCCTAACCC
anti-teloC: GGGTTAGGGTTAGGGTTAGGG

### Immunolocalization and quantitative analyses for mouse and human paraffin-embedded tissues

Four-micrometers-thick human and mouse tissue sections were deparaffinized, rehydrated and unmasked using Novocastra Epitope Retrieval Solutions pH8 in thermostatic bath at 98°C for 30 minutes. Subsequently, the sections were brought to room temperature and washed in PBS. After neutralization of the endogenous peroxidase with 3% H2O2 and Fc blocking by a specific protein block (Leica Novocastra), the samples were incubated with the primary antibodies.

IHC staining was revealed using Novolink Polymer Detection Systems (Leica Novocastra) or IgG (H#L) specific secondary antibodies (Life Technologies, 1:500) and DAB (3,3’-Diaminobenzidine, Leica Novocastra) as substrate chromogen.

For multiple-marker immunostainings, sections were incubated with ACE2 and TTF-1 primary antibodies and the binding of the primary antibodies to their respective antigenic substrates was revealed by made-specific secondary antibodies conjugated with Alexa-488 (Life Technologies, 1:250) and Alexa-568 (Life Technologies, 1:300) fluorochromes. In order to multiplex antibodies raised in the same species (ACE2 and Prosurfactant Protein C), Opal Multiplex IHC kit was developed. After deparaffinization, antigen retrieval in pH8 buffer was brought to a boil at 100% power, followed by 20% power for 15 minutes using microwave technology (MWT). Sections were treated with blocking buffer for 10 minutes at room temperature before primary antibody incubation. Slides were then incubated with Polymeric horseradish peroxidase-conjugated (HRP) secondary antibody for 10 minutes and the signal was visualized using Opal 520 fluorophore-conjugated tyramide signal amplification (TSA) at 1:100 dilution. The HRP catalyze covalent deposition of fluorophores around the marker of interest. The slides were again processed with the microwave treatment to strip primary/secondary antibody complex and allow the next antigen-antibody staining. Another round of staining was performed with the second primary antibody incubation, followed by Polymeric horseradish peroxidase-conjugated (HRP) secondary antibody and Opal 620 fluorophore-conjugated tyramide signal amplification (TSA) at 1:100 dilution for signal visualization. Finally, slides were again microwaved in antigen retrieval buffer and nuclei were subsequently visualized with DAPI (4’,6-diamidin-2-fenilindolo).

Slides were analyzed under a Zeiss Axioscope A1 microscope equipped with four fluorescence channels widefield IF. Microphotographs were collected using a Zeiss Axiocam 503 Color digital camera with the Zen 2.0 Software (Zeiss).

Quantitative analyses of immunohistochemical stainings were performed by calculating the average percentage of positive signals in five non-overlapping fields at medium-power magnification (x200) using the Positive Pixel Count v9 Leica Software Image Analysis.

### Immunofluorescence for OCT embedded mouse tissues

A measure of 4-mm tissue sections were fixed for 10 min in 4% PFA, incubated in blocking solution (2% BSA, 0.1% Tween in PBS) for 1 hour at RT. Then sections were incubated for 1 h at RT with primary antibodies, washed in blocking solution and incubated for 1 h at RT with secondary antibody. Nuclei were stained with DAPI (1 mg ml^−1^). Samples were mounted with glycerol solution. Image acquisition was performed in a Leica TCS SP5 confocal microscope. The detection parameters were set in the control samples and were kept constant across specimens.

### Immunofluorescence for cultured cells

Cells were fixed with 4% PFA. After incubation with blocking solution, cells were stained with primary antibody for 1 h at RT, washed and incubated with secondary antibodies for 40 min at RT. Nuclei were stained with 4,6-diamidino-2-pheny-lindole (DAPI; 1 mg ml^−1^). Samples were mounted in mowiol. Image acquisition was performed in a Leica AOBS SP2 confocal microscope. The detection parameters were set in the control samples and were kept constant across specimens.

### ImmunoFISH for cultured cells

After the immunofluorescence protocol, cells were treated as described (https://bioprotocol.org/e999). Briefly, after incubation with the secondary antibodies, cells were washed, fixed with PFA 4% + triton 0.1% for 10 min RT, incubated with glycine 10 mM for 30 min RT, washed, denatured at 80°C for 5 min and incubated with Cy5-conjugated TelC PNA probe (F1003 - Panagene). After washes, nuclei were stained with 4,6-diamidino-2-pheny-lindole (DAPI; 1 mg ml^−1^). Samples were mounted in mowiol. Image acquisition was performed in a Leica AOBS SP2 confocal microscope. The detection parameters were set in the control samples and were kept constant across specimens.

### Antibodies

The following primary antibodies were adopted for IHC and IF on mouse and human tissues: rabbit polyclonal ACE2 (1:500 pH8, ab15348, Abcam), rabbit polyclonal Prosurfactant Protein C (1:200 pH9, AB3786, Merck Millipore), mouse monoclonal TTF-1 (clone SPT24, 1:100 pH6, NCL-L-TTF-1, Leica Novocastra), rabbit polyclonal anti-53BP1 (1:1000, sc-22760, Santa Cruz), mouse monoclonal anti-phospho-Histone H2A.X (1:500, 05-636, Sigma-Aldrich).

### RNA extraction from mouse tissues

To isolate RNA, 20–30 mg lung tissue was homogenized in Trizol (Life Technologies) with Tissue Lyzer II (Qiagen) and processed with RNeasy kit (Qiagen) according to the manufacturer’s specifications. To increase the purity of the RNA extracted a convenient on-column DNase (Qiagen) treatment was performed to remove the residual amounts of DNA.

### RNA extraction from cultured cells

For MEFs, HeLa, and BJ cells total RNA was extracted using Maxwell® RSC Instrument, with Maxwell® RSC simplyRNA Tissue Kit (AS134, Promega), following manufacturer’s instructions. NanoVue™ Plus Spectrophotometer (GE Healthcare) was used to detect RNA quantity and purity. RNA purity was ascertained via NanoVue 260/280 and 260/230 ratios.

For HBEC, total RNA was extracted using QIAGEN RNeasy Plus Kit (74034. Qiagen) following manufacturer’s instructions. NanoDrop spectrophotometer was used to detect RNA quantity and purity. RNA purity was ascertained via NanoDrop 260/280 and 260/230 ratios.

Reverse Transcription Quantitative PCR (RT-qPCR)

For MEFs, HeLa and BJ 1-2 μg of total cell RNA was reverse transcribed into cDNA using SuperScript VILO cDNA Synthesis Kit (11754050, ThermoFisher). A volume corresponding to 25-50 ng of cDNA was used for each RT-qPCR reaction using Roche LightCycler 480 SYBR Green I Master (04707516001, Roche) sequence detection system. Each reaction was performed in triplicate.

Primers sequences (5’–3’ orientation) were:

Total ACE2 as described in (Ma *et al*, 2020)
  Mouse ACE2_Forward: TCCATTGGTCTTCTGCCATCC
  Mouse ACE2_Reverse: AACGATCTCCCGCTTCATCTC
  Human ACE2_Forward: TCCATTGGTCTTCTGCCATCC
  Human ACE2_Reverse: AACGATCTCCCGCTTCATCTC
Specific primers for ACE2 isoforms as described in (Blume *et al*, 2021)
  ACE2_Short_Fw: GTGAGAGCCTTAGGTTGGATTC
  ACE2_Short_Rv: TAAGGATCCTCCCTCCTTTGT
  ACE2_Long_Fw: CAAGAGCAAACGGTTGAACAC
  ACE2_Long_Rv: CCAGAGCCTCTCATTGTAGTCT
  ACE2_Total_Fw: TGGGACTCTGCCATTTACTTAC
  ACE2_Total_Rv: CCCAACTATCTCTCGCTTCATC

Ribosomal protein large P0 (Rplp0) RNA was used as control transcript for normalization:

Mouse/Human Rplp0_Forward: TTCATTGTGGGAGCAGAC
Mouse/Human Rplp0_Reverse: CAGCAGTTTCTCCAGAGC

### Digital droplet PCR

For HBEC, 200 ng of RNA was reverse transcribed into cDNA with iScript cDNA synthesis kit (1708890. Bio-Rad) following manufacturer’s instructions.

ddPCR of 20ng and 30ng cDNA were performed using QX 200 ddPCR EvaGreen supermix according to standard protocols (Ludlow *et al*, 2014).

Primer sequences (5’-3’ orientation) were:

ACE2 Forward Primer: TGTTGGGGAAATCATGTCACT
ACE2 Reverse Primer: GAGCAGGAAGTTTATTTCTGTTTCA

Amplicon size was 112 nt. Primers were designed, via primer-blast (Ye *et al*, 2012) and Roche assay design center, to be intron spanning and specific to ACE2 mRNA.

### Strand-specific real-time quantitative PCR

Detection of tdilncRNAs was performed as previously described (Rossiello *et al*, 2017), with some modifications. Briefly, RNA samples were treated with DNase I (Thermo Scientific) at 37 °C for 1 h. Next, 1 μg of total RNA was reverse transcribed using the Superscript First Strand cDNA synthesis kit (Invitrogen) with strand-specific primers. cDNA was passed on a MicroSpin™ G-50 columns (Cytiva) and qPCR was performed using SYBR Green I Master Mix (Roche). A volume of cDNA corresponding to 20 ng of initial RNA was used. Each reaction was performed in triplicate. Rplp0 was used as a control gene for normalization.

Primer sequences (5−3′ orientation):

Rplp0 Fw TTCATTGTGGGAGCAGAC
Rplp0 Rv CAGCAGTTTCTCCAGAGC
teloC Rv CCCTAACCCTAACCCTAA
teloG Rv GGGTTAGGGTTAGGGTTA
telo Fw CGGTTTGTTTGGGTTTGGGTTTGGGTTTGGGTTTGGGTT
telo Rv GGCTTGCCTTACCCTTACCCTTACCC TTACCCTTACCCT

### Luciferase assay

HeLa shTRF2 cells were transfected with ACE2(−1119)-luc plasmid, a gift from Gerhart Ryffel (Addgene plasmid # 31110; http://n2t.net/addgene:31110; RRID:Addgene_31110), 5 days after doxycycline treatment to induce shTRF2 expression or 1 day after the ionizing radiations treatment. Luciferase activity was measured 24 h post-transfection using the Luciferase Assay System (E4550, Promega) accordingly to the manufacturer instructions. Luciferase activity was normalized on total protein quantities.

### Single cell transcriptomic analyses

Single-cell transcriptomic data from ageing tissues in the mouse lung were downloaded from Tabula Muris Senis (https://cellxgene.cziscience.com/collections/0b9d8a04-bb9d-44da-aa27705bb65b54eb (Almanzar *et al*, 2020)). Seurat (v4.0.1, (Hao *et al*, 2020)) was used for downstream analyses.

### In silico transcription factor binding sites analysis

The tool Pscan (Zambelli *et al*, 2009) was used to look for over-represented transcription factor binding site motifs in nucleotide sequences. TFBS motifs for the human Ace2 gene (refseq ID: NM_021804) were searched around −450 +50 of the TSS, using the Jaspar 2018_NR matrix as a descriptor.

The EnrichR tool (Kuleshov *et al*, 2016) was used to perform a gene set enrichment analysis of the resulting Top100 significantly over-represented transcription factors.

### Statistical analyses

Results are shown as mean ± s.e.m. or s.d. or percentages ±95% confidence interval as indicated. P value was calculated by the indicated statistical tests, using Prism software. P values for single cell data analyses were calculated using the indicated statistical tests using the R software environment or with the default tests of the tools used.

## Acknowledgements

We thank Joachim Lingner and Marco Giorgio for sharing reagents and all F.d’A.d.F. laboratory members for discussions.

S.S. is supported by Fondazione Umberto Veronesi (FUV) and was previously supported by SIPOD 2 (Structural International Post Doc Programm 2) — the People Programme (Marie Curie Actions) of the European Union’s Seventh Framework Programme FP7 under grant agreement n.600399. F.d’A.d.F laboratory is supported by: ERC advanced grant (TELORNAGING – 835103); AIRC-IG (21762); Telethon (GGP17111); AIRC 5X1000 (21091); ERC PoC grant (FIREQUENCER – 875139); Progetti di Ricerca di Interesse Nazionale (PRIN) 2015 “ATR and ATM-mediated control of chromosome integrity and cell plasticity”; Progetti di Ricerca di Interesse Nazionale (PRIN) 2017 “RNA and genome Instability”; Progetto AriSLA 2021 “DDR & ALS”; POR FESR 2014-2020 Regione Lombardia (InterSLA project); FRRB - Fondazione Regionale per la Ricerca Biomedica - under the frame of EJP RD, the European Joint Programme on Rare Diseases with funding from the European Union’s Horizon 2020 research and innovation programme under the EJP RD COFUND-EJP N° 825575.

---

**Figure EV1:**
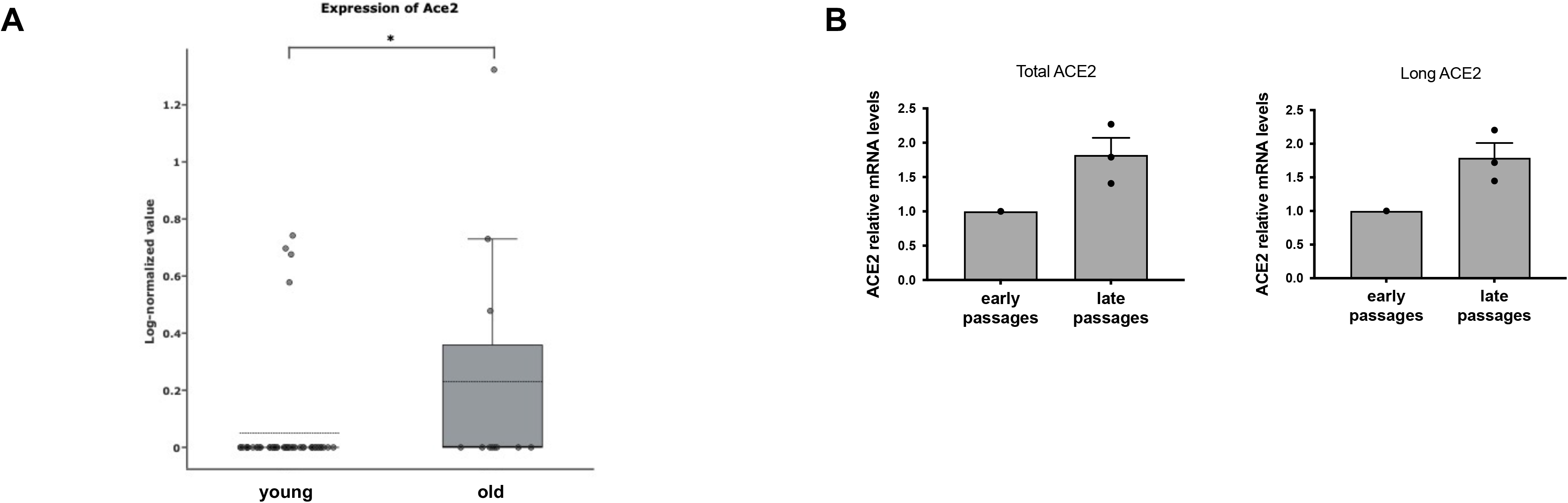
Supporting information. **A** Boxplot of *Ace2* expression levels in type II pneumocytes of young (3 months) and old (30 months) mice from single-cell transcriptomic data of aging tissues in the mouse lung from Tabula Muris Senis. *P<0.05, one-tailed Wilcoxon test. **B** RT-qPCR detection of *ACE2* mRNA expression levels in early passages (PD 35) and late passages (PD 61-64) BJ, using primers specific for the total or the long isoform (n = 1-3 replicates per group).

**Figure EV2.**
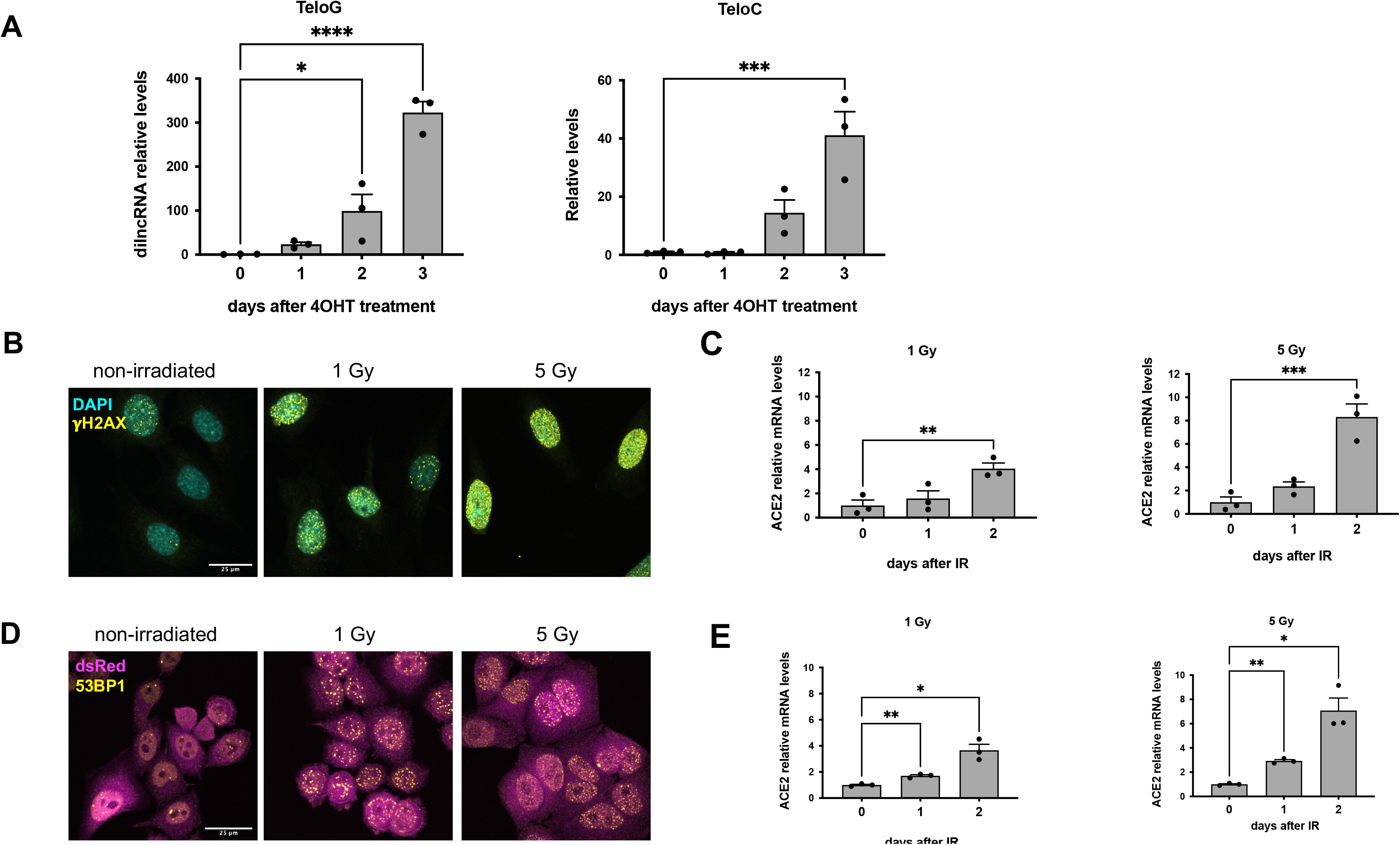
Supporting information. **A** RT-qPCR detection of telomeric dilncRNA expression levels in MEFs Trf2^F/F^ following 4OHT treatment (n=3 independent experiments). Error bars represent the s.e.m. *P<0.05, ***P<0.001,****P<0.0001. **B** Immunofluorescence showing γH2AX foci 6 hours after the indicated dose of ionizing radiation in MEFs Trf2^F/F^. Scale bar, 25 μm. **C** RT-qPCR detection of *Ace2* expression levels in MEFs Trf2^F/F^ treated as in B (n=3 independent experiments). Error bars represent the s.e.m. **P<0.01, ***P<0.001. **D** Immunofluorescence showing 53BP1 foci 6 hours after the indicated dose of ionizing radiation in HeLa shTF2. **E** RT-qPCR detection of *ACE2* expression levels in HeLa shTRF2 treated as in D (n=3 independent experiments). Error bars represent the s.e.m. **P<0.01, ***P<0.001.

